# Mechanisms of DNA repair have evolved to minimise the probability of nonsense mutations

**DOI:** 10.1101/2021.06.22.449367

**Authors:** George Powell, Helen S Long, Thomas M M Versluys, Ann-Marie Mallon, Cecilia M Lindgren, Michelle M Simon

**Author notes:** Denotes corresponding author.

## Abstract

Variation in sequence mutability has important implications for evolutionary models and predicting disease occurrence, and is driven in part by evolutionary divergence in mechanisms of DNA repair. The aim of this study was twofold: first, to assess the effect of local sequence context on substitution rates in the mouse lineage; second, to investigate the relationship between sequence mutability and selection. We show that the 7-mer context (i.e three bases either side of the base of interest) explains more variation in substitution rates between chromosomes in the mouse lineage than either the 3-mer, 5-mer, or 9-mer contexts. Furthermore, we also show that 7-mer substitutions with the potential to cause nonsense mutations when they occur in translated sequences occur at a lower rate across the genome than 7-mer substitutions with the potential to cause synonymous mutations. We propose that mechanisms of DNA repair have evolved to prioritise substitutions that are more likely to be deleterious to fitness.

## INTRODUCTION

Germline mutations are the basis of evolution and cause of inherited Mendelian and complex disease. Most mutations are either neutral or deleterious to fitness, and therefore organisms have evolved mechanisms of repairing DNA to maintain sequence fidelity (Harris and Pritchard 2017). These mechanisms are complex, involving hundreds of genes in eukaryotes (Lindahl and Wood 1999; Ségurel, Wyman, and Przeworski 2014), and are known to diverge between lineages over evolutionary time, resulting in variation in sequence mutability (Harris 2015; Kumar and Subramanian 2002; Lindsay et al. 2019). Quantifying this variation in sequence mutability is important for parameterising evolutionary models, interpreting patterns of variation, and identifying mutational signatures and pathogenic variants (Aggarwala and Voight 2016; Samocha et al. 2014; Karczewski et al. 2020; Helleday et al. 2008).

Variation in sequence mutability is influenced by local sequence context. For example, substitution rates are affected by a base’s k-mer context, which describes the composition of its flanking nucleotides (Harris and Pritchard 2017; Aikens, Johnson, and Voight 2019; Schroeder et al. 2016). In human populations, the 7-mer context (ie three bases either side of the base of interest) has been shown to account for more variation in substitution rates than either the 3-mer or 5-mer context (Aggarwala and Voight 2016). Variation in mutability between k-mers is partly driven by the chemical properties of bases and their epigenetic state (Ségurel, Wyman, and Przeworski 2014; Gonzalez-Perez, Sabarinathan, and Lopez-Bigas 2019; Makova and Hardison 2015). For example, the higher mutability of C-G relative to A-T base pairs is partly attributed to the difference in their number of connecting hydrogen bonds ((Petruska and Goodman 1985; Hardison et al. 2003); (Ségurel, Wyman, and Przeworski 2014)), and C > T substitutions are enriched amongst CG dinucleotides due to methylation and spontaneous cytosine deamination (Shen, Rideout, and Jones 1994; Duncan and Miller 1980). However, variation in k-mer mutability is also affected by mechanisms of DNA repair, which can diverge over evolutionary time. For example, the TCC > TTC substitution rate has increased by 50% in the human European lineage since Europeans began diverging from Asians as a result of changes to DNA repair mechanisms (Harris 2015). The cause of these changes remains unclear, but they are tentatively linked to differences in UV exposure. TCC > TTC is the substitution most frequently induced by UV and present in melanoma skin cancers, and it is speculated that decreased exposure to UV in the European lineage may have reduced the selection pressure on mechanisms for repairing TCC > TTC substitutions, resulting in their relative increase in rate of occurrence (Harris 2015). Nevertheless, determining how selection may have affected sequence mutability through the evolution of mechanisms of DNA repair remains poorly understood and is difficult to test empirically.

Here, we assess the effect of local sequence context on substitution rates across the mouse genome using genetic variation between mice sampled from wild populations (Harr et al. 2016). Mice are the most commonly used mammalian model organism for biomedical research, and offer important insights into human biology (Brown et al. 2018). As mutation rates diverge over evolutionary time, quantifying sequence mutability in the mouse lineage can be used to parametrize mouse-specific models for inferring functional genomic regions under negative selection, and identify mutational signatures and pathogenic variants (Karczewski et al. 2020; Helleday et al. 2008; di Iulio et al. 2018). Our research has two main aims. First, we assess the contribution of flanking nucleotides to the ability to predict substitution mutations in the mouse lineage. We show that the 7-mer context explains more variation in substitution occurrence between chromosomes than either the 3-mer, 5-mer, or 9-mer contexts. Second, we assess the relationship between sequence mutability and selection. Specifically, we hypothesise that substitutions that are more likely to be deleterious to fitness have lower base rates of occurrence. In support of this hypothesis, we show that 7-mer substitutions with the potential to cause nonsense mutations when they occur in translated sequences occur at a lower rate across the genome than 7-mer substitutions with the potential to cause synonymous mutations. We propose that this variation in mutability is the result of selection on mechanisms of DNA repair.

## RESULTS

### The effect of local sequence context on substitution rates

We assessed the effect of local sequence context on the ability to predict substitution mutations in the mouse lineage. We hypothesised that considering a greater number of flanking nucleotides either side of the base of interest (ie expanding the sequence context) would improve the ability to predict substitutions across the mouse genome.

We considered six different k-mer models to account for local sequence context (supplementary table 1): the 1-mer, 3-mer, 5-mer, 7-mer, 9-mer and A-CG models. The 1-mer model is the simplest model and does not consider the local sequence context of the base of interest. It describes 12 substitution rates, as each of the four bases has three possible substitutions. The 3-mer model considers one flanking nucleotide either side of the base of interest, making a 3-mer. It describes 192 substitution rates as there are 64 possible 3-mers and the middle base in each 3-mer has three possible substitutions. The 5-mer, 7-mer, and 9-mer models consider 2, 3, and 4 flanking nucleotides either side of the base of interest, respectively. The number of substitution rates that models describe increases exponentially as the number of flanking nucleotides expands, with the 9-mer model describing 786,432 substitution rates as there are 262,144 possible 9-mers and the middle base in each 9-mer has three possible substitutions. Substitution probabilities are known to be largely driven by the presence of CG dinucleotides, which have a high mutation rate from cytosine to thymine (M. Li and Chen 2011). We therefore expanded our simple 1-mer model to consider CG context, and named this the A-CG model. The A-CG model describes 18 substitution rates as there are 6 possible contexts (A, T, C (non-CG), C (CG), G (non-CG), and G (CG)), each with three possible substitutions. For each k-mer model we additionally calculated the probability of any mutation as the sum of each potential substitution for each k-mer.

To estimate substitution rates we considered autosomal single nucleotide variants (SNVs) from populations of wild mice (*Mus musculus* sp. and *Mus spretus*) (Harr et al. 2016), and inferred the ancestral state of each allele from alignment with the *Mus caroli* reference genome. We filtered out 75.3% of the mouse genome, including genomic regions with low sequence coverage, repetitive elements, and regions with functional annotation such as coding and regulatory features that are likely to be under selection (see methods for more detail). We assessed the accuracy of each k-mer model to predict substitution occurrence using train/test resampling by chromosome. In brief, fifteen autosomes were randomly selected to calculate k-mer probabilities of substitution and the remaining four autosomes were used to test predictive accuracy using the absolute error between the observed substitution rates (test autosomes) and expected substitution rates (training autosomes). This was repeated 100 times to provide the mean absolute error (MAE) and 95% confidence intervals. Each ancestral k-mer and k-mer change has a complement on the reverse strand. For example, the ancestral 3-mer TAG on the forward strand must have the complementary k-mer CTA on the reverse strand, and a TAG > TTG substitution must result in a complementary CTA > CAA substitution. We therefore totaled the counts of each forward and reverse k-mer and k-mer change to calculate substitution probabilities, and report the substitution probabilities for adenine and cytosine only. This results in six substitution classes under consideration (noted as A>C, A>G, A>T, C>A, C>G, and C>T), plus the occurrence of any adenine or cytosine substitution (noted as A>* and C>*).

Of the k-mer models considered, the 7-mer context provides the greatest increase in predictive accuracy relative to the 1-mer and A-CG models for each substitution class in the mouse lineage (figure 1). The 7-mer model reduces mean absolute error (MAE) by 73.7% ± 0.3% relative to the 1-mer model for cytosine substitutions (C > *). This is largely driven by the presence of CG dinucleotides and the frequency of CG > TG substitutions, with the A-CG model reducing MAE by 67.0% ± 0.3% for cytosine substitutions relative to the 1-mer model (figure 1). The 7-mer model, however, reduces MAE for cytosine substitutions by 20.5%±0.9% relative to the A-CG model, indicating the additional nucleotides either side of a base of interest affect its substitution rate. This pattern is consistent for adenine substitutions (A > *), with the 7-mer model reducing MAE by 34.5% ±0.7% relative to the 1-mer model.

**Figure 1.**
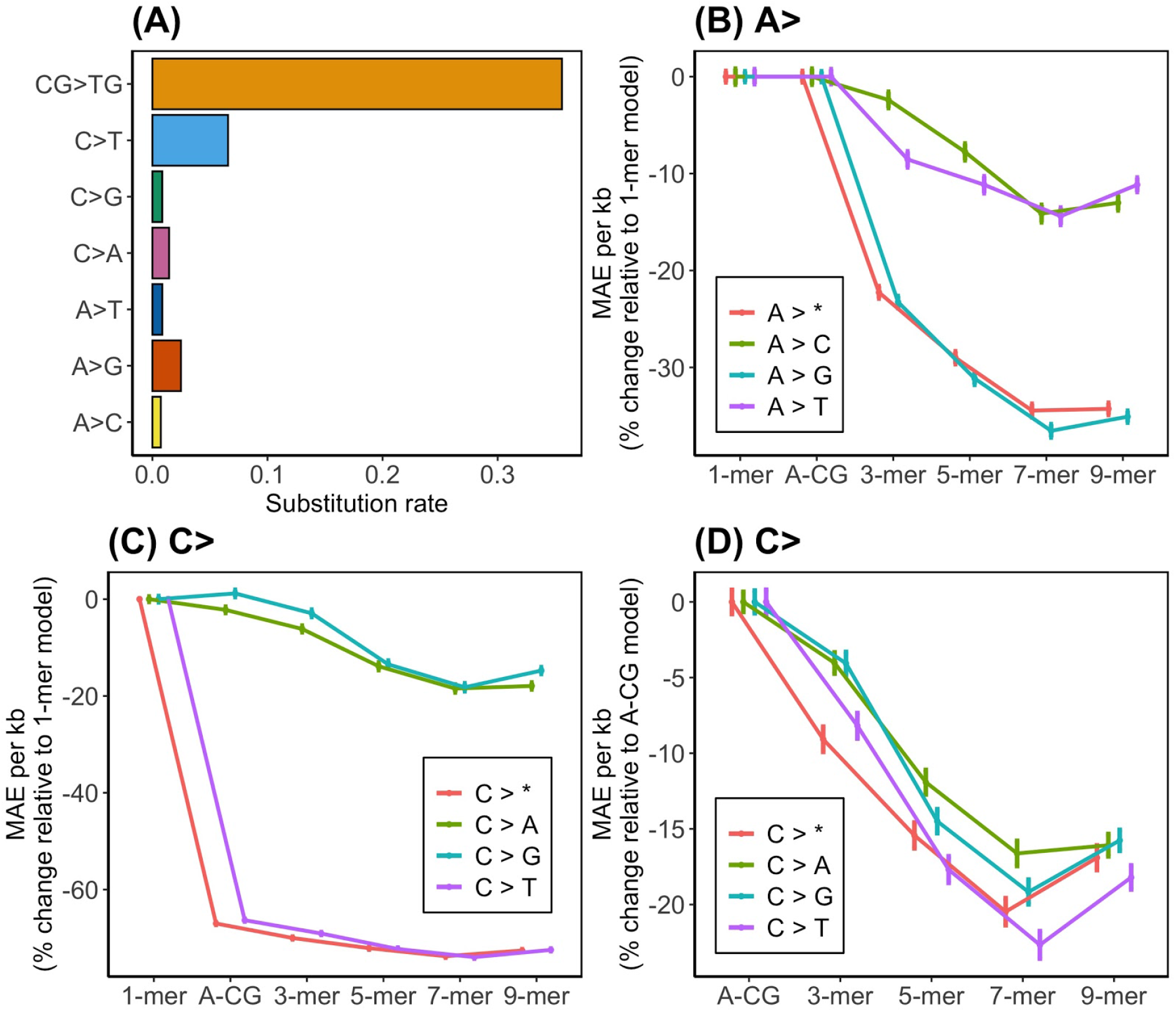
A) 1-mer substitution rates by class. C > T substitutions occur at the greatest rate, largely driven by the mutability of CG > TG dinucleotides. B) Mean absolute error (MAE) per kb relative to the 1-mer model for predicting adenine substitutions. The 7-mer model provides the greatest improvement in MAE relative to the 1-mer model for all substitution classes, with the greatest improvement for A > G substitutions (36.5% ±0.7%). C) Mean absolute error (MAE) per kb relative to the 1-mer model for predicting cytosine substitutions. The A-CG model decreases MAE by 67.0% ±0.3% for C > T substitutions relative to the 1-mer model by considering the CG context. D) Mean absolute error (MAE) per kb relative to the A-CG model for predicting cytosine substitutions. The 7-mer model provides the greatest improvement in MAE relative to the A-CG model for all substitution classes, with the greatest improvement (22.7% ±0.9%) shown for C > T substitutions. Any potential substitution is denoted with a *.

The 9-mer model explains less variance than the 7-mer model, suggesting that a further increase in bases does not provide further information to account for substitution occurrence (figure 1). However, it is possible that due to the increased number of parameters in the 9-mer model there is insufficient statistical power to detect the improvement. To test this, we simulated a null dataset where the counts of each 9-mer on each chromosome is equal to the observed counts, and the probability of substitution is dependent on the 9-mer context (see methods). Given this simulated dataset, we found the MAE was lower for the 9-mer than the 7-mer model (supplementary figure 2) for all substitution classes except C > T. This suggests we were not lacking statistical power to detect an effect of the simulated magnitude for each substitution class other than C > T. The distributions of k-mer and k-mer substitution counts are provided in supplementary table 8.

Genomic regions with high GC content have higher mutation rates (Ségurel, Wyman, and Przeworski 2014). This is hypothesised to result from decreased efficiency of exonuclease activity in separating C-G base pairs relative to A-T in order to excise erroneous nucleotides, because A-T base pairs are bound by two hydrogen bonds whereas C-G base pairs are bound by three (Petruska and Goodman 1985; Hardison et al. 2003). We fit a generalised linear model to predict the number of cytosine or guanine bases in the 7-mer as a function of its substitution rate, and found a significant positive relationship (estimate = 0.82±0.03, z = 25.7, p = 2.66e-167) (supplementary figure 4). To test whether this relationship is inflated by CG deamination we fit another model which excluded all CG>TG substitutions, and found a significant positive relationship remained (estimate = 1.90±0.12, z = 15.7, p = 1.18e-55). These results highlight that the relationship between GC content and mutability persists at a finer scale, and are consistent with the hypothesis that higher substitution rates are caused in part by decreased efficiency of exonuclease activity in separating C-G base pairs relative to A-T in order to excise erroneous nucleotides.

### Methylation and CG dinucleotide substitution rates

Methylation state affects the probability of substitutions within CG dinucleotides (figure 2). CG dinucleotides are highly mutable across the genome, particularly C > T substitutions resulting from deamination at methylated sites (Shen, Rideout, and Jones 1994; Duncan and Miller 1980). We quantified the effect of methylation state on the CG mutability in the mouse lineage. We considered all CG dinucleotides across unmasked regions of the genome, and inferred methylation status in the germline from bisulfite sequencing in embryonic stem cells, which show a similar pattern of methylation to sperm cells in mice (Popp et al. 2010). In total, 72.9% of CGs are classified as methylated, 6.9% as unmethylated, and 20.2% were unclassified due to insufficient coverage. The C > T substitution rate in methylated CG dinucleotides is 0.37 compared with 0.04 in unmethylated CGs, an 8.3 fold increase. By comparison, the C > T substitution rate is 0.35 across all CGs independent of methylation status. This is closer to the substitution rate for methylated CGs as there is a higher proportion of methylated CGs relative to unmethylated CGs in our sample. Accounting for methylation state improves prediction accuracy for all CG substitution classes, with the effect greatest for C > T substitutions (figure 2). MAE is reduced by 82.5%±0.2.5% when predicting the occurrence of C > T substitutions across methylated CGs, and 99.7%±0.1% across unmethylated CGs, relative to a null model where methylation status is not considered. The improvement in MAE is greater for unmethylated CGs relative to methylated CGs because the null substitution rate (calculated across all CGs independent of methylation status) has a higher proportion of methylated CGs relative to unmethylated CGs. The improvement in MAE by considering methylation status is greater than consideration of the 7-mer context alone, which improves prediction accuracy of C > T substitutions by 22.7%±0.9% relative to the A-CG model, further highlighting the importance of methylation state to predicting C > T substitution occurence.

**Figure 2.**
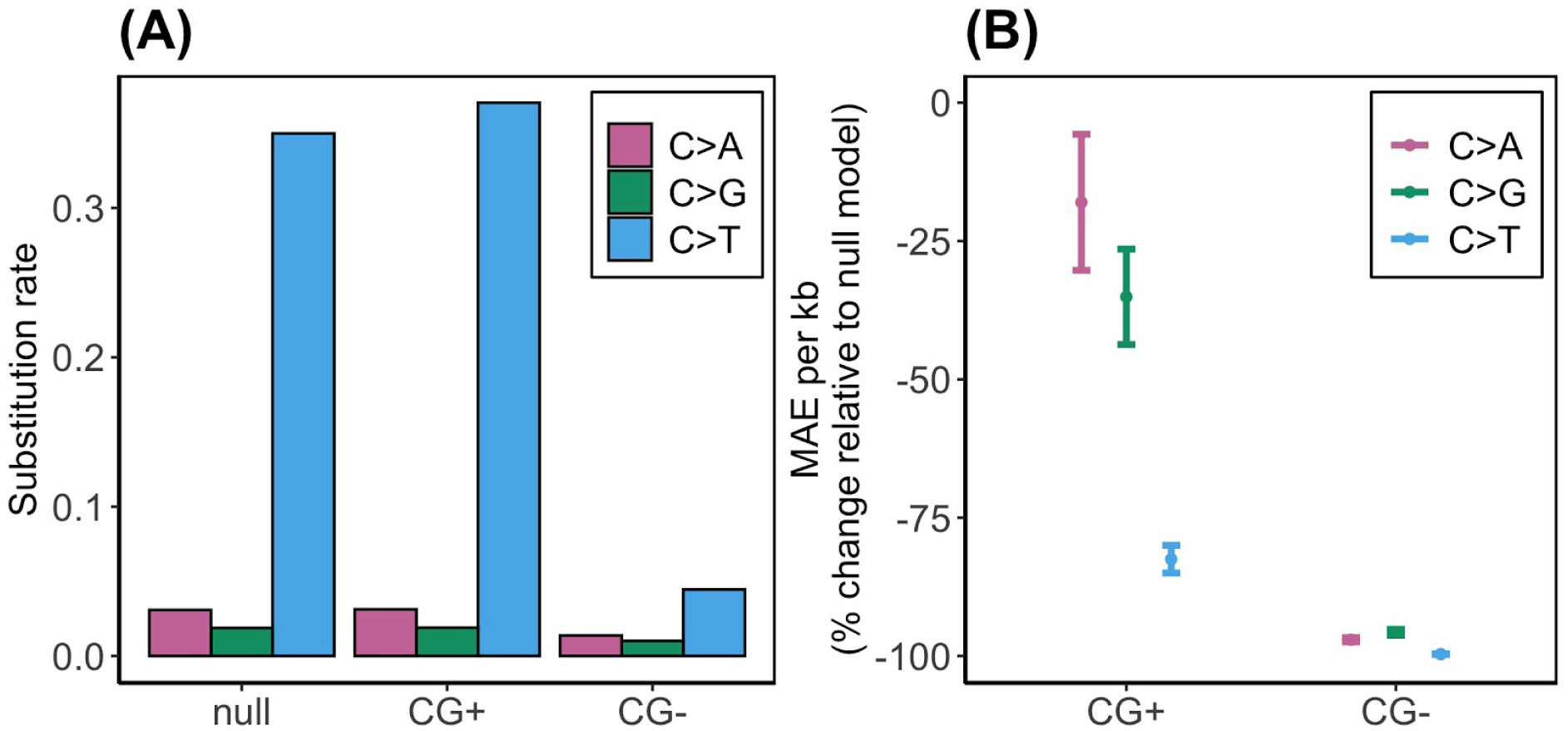
A) Substitution rates of methylated (CG+), unmethylated (CG−), and all (null) dinucleotides by class. Methylated CG dinucleotides have a C > T substitution rate of 0.37 compared with 0.04 for unmethylated CG dinucleotides. In total, we classified 72.9% of CG dinucleotides across the unmasked genome as methylated, 6.9% are unmethylated, and 20.2% are unclassified because of low coverage. B) Mean absolute error (MAE) per kb relative to the null model, in which methylation status was not considered, for methylated (CG+) and unmethylated (CG−) dinucleotides. Considering methylation status reduces MEA for each substitution class. The greatest decreases are for C > T substitutions, with a MAE decrease of −82.5%±0.2.5% and −99.7%±0.1% for methylated and unmethylated CG dinucleotides, respectively.

### Mutational effect and probability of occurrence

Mechanisms for DNA repair affect rates of substitution and can diverge over evolutionary time (Harris 2015; Harris and Pritchard 2017). We hypothesise that k-mer substitutions that are more likely to be deleterious to fitness will occur at lower frequencies across the genome resulting from selection on mechanisms of DNA repair. For example, if AAC > ACC substitutions are more likely to be deleterious to fitness than AAG > ACG, there would be a stronger selection pressure on DNA repair mechanisms to repair AAC > ACC compared with AAG > ACG, resulting in a relative decrease of AAC > ACC substitutions. To assess this we looked at the potential of 7-mer substitutions to cause nonsense and synonymous mutations if they occur in translated regions of the genome, as nonsense mutations tend to be under strong negative selection relative to synonymous mutations. Specifically, we tested three hypotheses. First, 7-mer substitutions with the potential to cause nonsense mutations if they occur in translated sequences have lower substitution rates across the genome than 7-mers substitutions with the potential to cause synonymous mutations. Second, the number of potential nonsynonymous mutations for each 7-mer substitution is negatively correlated with its substitution rate, and conversely the number of potential synonymous mutations for each 7-mer substitution is positively correlated with the substitution rate. Finally, rates of 7-mer substitutions with the potential to cause nonsense mutations will be more conserved between species than rates of 7-mer substitution with the potential to cause synonymous mutations.

We first tested whether 7-mer substitutions with the potential to cause nonsense mutations if they occur in translated sequences have lower substitution rates across the genome (ie across translated and untranslated regions) than 7-mers substitutions with the potential to cause synonymous mutations. To test this hypothesis, we grouped 7-mer substitutions by their potential to cause nonsense, missense, or synonymous mutations if they occur in translated sequences (supplementary figure 3) and tested the difference in observed substitution rates between groups. Substitution rates were calculated from the unmasked regions of the genome as previously described and excluded translated sequences (see methods). Nonsense or missense mutations are more likely to be under negative selection than synonymous mutations, and we therefore expect 7-mer substitutions with the potential to cause nonsense or missense mutations in translated sequences to have lower substitution rates than 7-mers substitutions with the potential to cause synonymous mutations. 7-mer substitutions with the potential to cause nonsense mutations when in translated regions have significantly lower rates of occurrence across the genome than those with the potential to cause synonymous mutations (W = 124021674, p = 5.8e-124)(figure 3). When considered across all substitution classes, this relationship is largely driven by the inability of A > G substitutions, which have on average higher substitution rates than either A > C or A > T substitutions, to cause nonsense mutations (figure 3). Differences do, however, remain when controlling for substitution class (ie A>C, A>G, A>T, C>A, C>G, and C>T) (supplementary table 9). For each substitution class, 7-mer substitutions with the probability of causing nonsense mutations have significantly lower probabilities of occurrence than 7-mer substitutions with the probability of causing synonymous mutations (supplementary table 9) (p < 0.001). There are more potential missense substitutions (n = 415) than nonsense (n = 23) or synonymous (n = 138), and all 7-mer substitutions (n = 24576) have the potential to cause at least one missense mutation. Potential to cause a missense substitution therefore describes the distribution of all 7-mer substitution rates.

**Figure 3.**
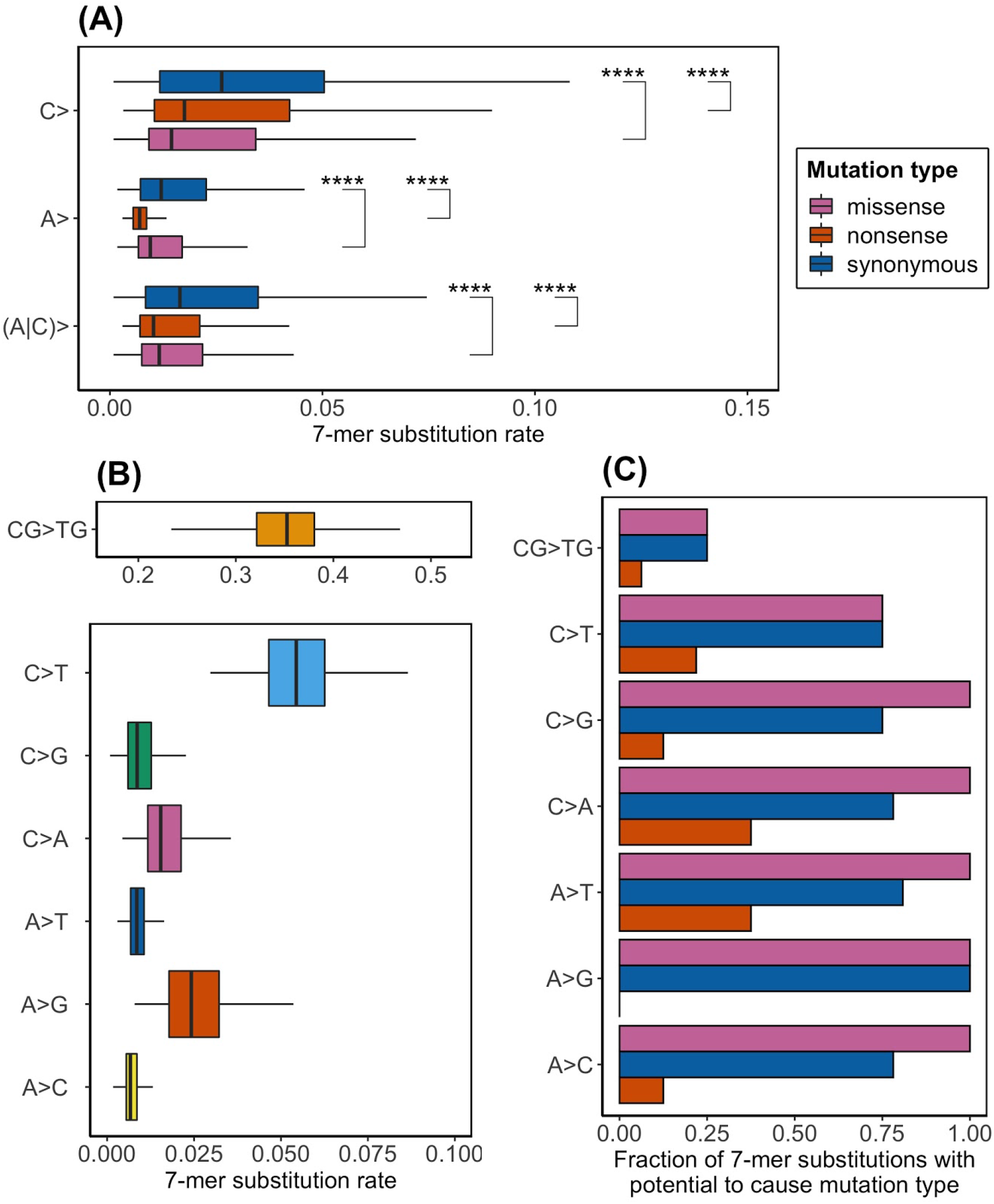
7-mer substitutions with the potential to cause nonsense and missense mutations when they occur in translated sequences have significantly lower probabilities of occurrence than 7-mer substitutions with the potential to cause synonymous mutations (A). This is largely driven by the inability of A > G substitutions, which have a higher substitution rate (B), to cause nonsense mutations (C). A) Distributions of substitution rates for 7-mers with the potential to cause synonymous, missense, and nonsense variation. “A > *” and “C > *” denote all possible adenine and cytosine substitutions and their complementary thymine and guanine substitutions, respectively. “(A|C) >” encompasses all possible substitutions. Differences between groups were assessed with Wilcoxon signed-rank tests, with *** denoting a p value less than 0.0001. Outliers, defined by being outside 1.5 times the interquartile range above the upper quartile and below the lower quartile, are not shown. B) 7-mer substitution rates by class. Outliers, defined by being outside 1.5 times the interquartile range above the upper quartile and below the lower quartile, are not shown. C) Fraction of 7-mer substitutions by type with the potential to cause nonsense, missense, and synonymous variants. All 7-mer substitutions have the potential to cause a missense mutation.

7-mer substitutions have the potential to cause multiple different mutation types when they occur in translated sequences (supplementary figure 3). For example, each 7-mer substitution can have six different consequences as they have the potential to change each of the three bases in a codon, and this applies to both the forward and reverse strands. We expect 7-mer substitutions with a higher probability of causing a nonsynonymous mutation to have lower rates of substitution, and 7-mer substitutions with a higher probability of causing a synonymous mutation to have higher rates of substitution. To test this we counted the number of potential synonymous and nonsynonymous mutations for each 7-mer substitution (supplementary figure 3). We grouped missense and nonsense substitutions to increase our power to detect a relationship, as the maximum number of nonsense mutations for any 7-mer substitution is two. We fit poisson generalised linear models to predict the number of each mutation type as a function of substitution rate for each 7-mer substitution. 7-mer substitutions with greater potential to cause nonsynonymous mutations have significantly lower substitution rates (coefficient = −0.67±0.05, z = −14.1, p = 4.2e-45), and 7-mer substitutions with greater potential to cause synonymous mutations have significantly higher substitution rates (coefficient = 0.85±0.06, z = 13.3, p = 1.5e-40).

Finally, we reasoned that if 7-mer substitution rates are under selection (or rather mechanisms of DNA repair mediating substitution rate) resulting from their potential to cause different mutation types in translated sequences, we would expect to see less variation in substitution rates between species for 7-mers with the potential to cause nonsense mutations relative to synonymous. This is because we expect negative selection to have minimised the substitution rates of potentially nonsense causing 7-mers in both lineages, whereas the substitution rates of potentially synonymous causing 7-mers if under weaker selection would be more likely to vary due to genetic drift. To test this we calculated rank correlation between human and mouse substitution rates for two groups: 7-mers with the potential to cause nonsense mutations and 7-mers with the potential to cause synonymous substitutions. For each substitution class, there is a significant correlation (p < 3.78e-126) and the correlation coefficient is greater for 7-mers with the potential to cause nonsense mutations than for 7-mers with the potential to cause synonymous substitutions (supplementary table 10). This highlights less variation in 7-mer substitution rates between species for those with the potential to cause nonsense mutations relative to synonymous.

## DISCUSSION

We have assessed the effect of local sequence context on substitution rates across the mouse genome, and the relationship between sequence mutability and selection. We have two main results: first, the three flanking nucleotides either side of the base of interest (its 7-mer context) explain the most variability in substitution rates between chromosomes in the mouse lineage; second, 7-mer substitutions that are more likely to be highly deleterious to fitness because they have the potential to cause nonsense mutations when they occur in translated sequences are less likely to occur across the genome.

The importance of 7-mer context to substitution occurrence has important implications for evolutionary models. In particular, models that aim to quantify the strength of selection from the relative enrichment or depletion of variation must control for regional differences in mutability (Samocha et al. 2014; di Iulio et al. 2018). Failure to do this may lead to false inferences of selection. For example, regions that are less mutable are likely to contain fewer variants, independent of selection. Our finding that 7-mer context improves predictive accuracy of substitution occurrence in mice is consistent with human studies reporting similar findings (Aggarwala and Voight 2016). This suggests that the causes of 7-mer context importance have been conserved between the species. These causes remain unclear, but they could involve both fundamental properties of DNA and DNA repair mechanisms. For example, sequence context may affect the ability of DNA polymerase to separate DNA in order to remove and replace an erroneous base. Erroneous bases that are surrounded by C or G nucleotides may be less likely to be corrected, as C-G nucleotides are joined by three hydrogen bonds and therefore require more energy to separate than A-T nucleotides, which are bound by two (Petruska and Goodman 1985; Hardison et al. 2003). The 7-mer context could account for the distance from an erroneous base by which DNA must be separated in order for repair. It remains to be determined whether the 7-mer context holds the same importance for non-mammalian species that are more distantly related to humans, such as drosophila.

Variation in sequence mutability between lineages captured by local sequence context suggests evolutionary divergence in mechanisms of DNA repair, with some lineages repairing specific substitutions more or less effectively than others (Ségurel, Wyman, and Przeworski 2014; Harris 2015; Kumar and Subramanian 2002). Evolutionary theory suggests the complexity of DNA proofreading and repair mechanisms in eukaryotes may limit the impact of selection on shaping the efficiency of individual genes in performing these processes (Sung et al. 2012; Lynch 2011). For example, if a variant changes the mutation rate in a given sequence context by a small amount then the effect on fitness may fall below the threshold at which selection is effective. Therefore, the fixation of the variant in the population would be dependent on genetic drift. This line of reasoning has led to the hypothesis that drift is the predominant cause of variation in mutability between species (Sung et al. 2012). We have shown, however, that 7-mer substitutions that are more likely to be deleterious to fitness (by having the potential to cause nonsense mutations in translated sequences) are less likely to occur across the genome. This pattern is consistent across each substitution class, and between humans and mice, indicating that when the strength of selection is strong enough, natural selection affects mutability. The genetic code that determines nonsense and synonymous mutations is universal to almost all species (Knight, Freeland, and Landweber 2001), but understanding whether selection drives further variation in mutability between lineages requires further research. For example, it would be interesting to identify k-mers known to be under lineage-specific negative selection, and determine whether this is reflected in their substitution rates.

## METHODS

### Data download and analysis software

We considered genetic variation between 67 mice (*Mus musculus* sp. and *Mus spretus*), sampled from 11 wild populations (Harr et al. 2016). Variant (vcf files) and coverage (bam files) information were downloaded from http://www.user.gwdg.de/~evolbio/evolgen/wildmouse/. The soft-masked mouse reference genome (GRCm38), *Mus musculus* to *Mus caroli* pairwise alignment (Herrero et al. 2016), GenCode exonic annotations (Harrow et al. 2012), and the Ensembl Regulatory Build (Zerbino et al. 2015) were all downloaded from Ensembl ftp (ftp://ftp.ensembl.org/pub/release-101//) (v101). GERP conserved regions were calculated by Ensembl across 111 mammalian species alignments, and were downloaded from Ensembl ftp (v101) (Herrero et al. 2016). Genome-wide methylation of CG dinucleotides in mouse embryonic stem cells was quantified by Stadler et al (2011) using bisulfite sequencing, and were downloaded from Ensembl ftp (http://ftp.ensembl.org/pub/data_files/mus_musculus/GRCm38/dna_methylation_feature/NPC_5mC_Publication_Stadler2011_PMID22170606/) (Stadler et al. 2011). Human 7-mer substitution rates were downloaded from the supplementary information of Aggarwala and Voight (2016).

All analyses were conducted using custom scripts written in R (v3.6.2) (R Core Team 2017) and bash, and are available on the following github repository: *https://github.com/powege/Thesis_workflow.git*.

### Filtering variants and genomic regions

We built a mask for the mouse genome (GRCm38) to filter low coverage regions, repetitive elements, and regions with functional annotation that are likely to be under selection. We filtered low coverage loci defined as having less than 10X coverage in more than 10% of individuals in the sample. Read depth at each genomic position for each mouse in the sample was calculated from the bam coverage files using samtools (H. Li et al. 2009). We filtered repetitive elements and ambiguous (N) bases by excluding all soft-masked regions classified using RepeatMasker by Ensembl (Zerbino et al. 2018). We also filtered regions likely to be under selection. These included functional regions annotated as either “promoter”, “promoter flanking”, “enhancer”, “CTCF binding site”, “TF binding site”, or “open chromatin” by the Ensembl Regulatory Build (v101) (Zerbino et al. 2015); all exonic regions annotated by GenCode (Harrow et al. 2012); and GERP evolutionarily conserved regions (Herrero et al. 2016). In total we masked 75.3% of the mouse genome.

We considered all 41,000,862 single nucleotide variants (SNVs) with “PASS” filter status across unmasked regions of the genome.

### Local sequence context models and probabilities of substitution

We considered six k-mer models for determining the effect of local sequence context on substitution probabilities (supplementary table 1): the 1-mer, 3-mer, 5-mer, 7-mer, 9-mer and A-CG models. The 1-mer model is the simplest model and does not consider the local sequence context of the base of interest. It has 12 parameters as each of the four bases has three possible substitutions. The 3-mer model considers one flanking nucleotide either side of the base of interest, making a 3-mer. It has 192 parameters as there are 64 possible 3-mers (4^3) and the middle base in each 3-mer has three possible substitutions. The 5-mer, 7-mer, and 9-mer models consider 2, 3, and 4 flanking nucleotides either side of the base of interest, respectively. The number of parameters increases exponentially as the number of flanking nucleotides considered expands, with the 9-mer model having 786,432 parameters as there are 262,144 possible 9-mers (4^9) and the middle base in each 9-mer has three possible substitutions. Substitution probabilities are largely driven by the presence of CG dinucleotides which have a high mutation rate from cytosine to thymine (M. Li and Chen 2011). We therefore expanded our simple 1-mer model to consider CG context, and named this the A-CG model. The A-CG model has 18 parameters as there are 6 possible contexts (A, T, C (non-CG), C (CG), G (non-CG), and G (CG)), each with three possible substitutions.

We inferred the mutational direction of each SNV (ie the ancestral and mutant states) using Ensembl’s alignment of the *Mus musculus* (GRCm38) and *Mus caroli* reference genomes. For each SNV, the ancestral state was assumed to be the reference allele unless the alternate allele was shared with *Mus caroli.*

We calculated the substitution probability for each k-mer by dividing the counts of each k-mer change by the counts of each ancestral k-mer across the unmasked genome (supplementary tables 2 to 7). Each ancestral k-mer and k-mer change has a complement on the reverse strand. For example, the ancestral 3-mer TAG on the forward strand must have the complementary k-mer CTA on the reverse strand, and a TAG > TTG substitution must result in a complementary CTA > CAA substitution. We therefore totaled the counts of each forward and reverse k-mer and k-mer change to calculate substitution probabilities, effectively halving the number of parameters for each model, and report the substitution probabilities for adenine and cytosine only.

### Methylation states and probabilities of substitution

Genome-wide methylation of CG dinucleotides in mouse embryonic stem cells was quantified by Stadler et al (2011) using bisulfite sequencing, and were downloaded from Ensembl (Stadler et al. 2011; Zerbino et al. 2018) (v101). Genome-wide methylation data is not available for mouse germline cells. However, similar patterns of methylation have been observed between mouse embryonic stem cells and sperm cells (Popp et al. 2010), and we therefore used mouse embryonic stem cells as a proxy for the germline.

We classified all CG dinucleotides across the unmasked mouse genome as methylated if they have a coverage greater than or equal to 5 and a percentage methylated greater than or equal to 60, and unmethylated if they have a coverage greater than or equal to 5 and a percentage methylated less than or equal to 20. We excluded CG dinucleotides with coverage less than 5, or a percentage methylated more than 20 and less than 60. The distribution of methylation percentages is shown in supplementary figure 1. In total we classified 72.9% of CG dinucleotides as methylated, 6.9% as unmethylated, and excluded 20.2% from our analysis.

We calculated the probability for cytosine substitutions across three CG dinucleotide groups: all CG dinucleotides, unmethylated CG dinucleotides, and methylated CG dinucleotides. For each unmasked CG dinucleotide we determined the substitution class by the alternate allele (ie C>A, C>G, C>T, G>A, G>C, G>T). We calculated probabilities of substitution for each CG group by dividing the counts of each substitution class in each CG group, by the number of CGs in each CG group. We then combined counts between complementary strands to leave three substitution probabilities: C>A, C>G, and C>T.

### Testing the prediction accuracy of substitution models

We used resampling by chromosome to test the prediction accuracy of local sequence context models. For each iteration, we split the data into training and testing datasets. Four autosomes were randomly selected to be the testing data, and the remaining fifteen autosomes were used to train the models. Due to differences in chromosome size, this meant that the testing data could range between 28.0% and 14.1% of the total dataset. Probabilities of substitution were calculated for each k-mer model using the training data, following the methodology described above. To minimise bias introduced by the composition of test chromosomes (ie some kmers appearing at a much higher frequency than others) we gave each 9-mer an equal weighting in calculating substitution occurence. This means the observed number of substitutions in the test data were calculated by totaling the observed substitution rates for each 9-mer. The absolute error was then calculated for each local sequence context model and substitution class (ie A>C, A>G, A>T, C>A, C>G, C>T) as the sum of the absolute difference between the observed (testing data) and predicted (training data) number of each substitution class for each 9-mer. Resampling was conducted 100 times and the mean absolute error and 95% confidence interval for each k-mer model and substitution class were calculated across the iterations.

We tested the accuracy to predict CG dinucleotide substitutions given different methylation states using resampling by chromosome, with four chromosomes randomly sampled for the test dataset in each iteration. Probabilities of substitution were calculated for each CG dinucleotide group in the training dataset following the methodology described above. Probabilities of substitution for methylated and unmethylated CG dinucleotides were calculated with the test dataset. The prediction accuracy of each CG group was determined as the difference between the observed (testing data) and predicted (training data) fraction for each substitution class.

### Power testing

The number of potential k-mers (ie the potential combinations of A, C, G, and T) increases exponentially with k-mer length (supplementary table 1). This reduces the average count for each k-mer and k-mer substitution in the sample (supplementary table 8), and therefore the statistical power to accurately determine probabilities of substitution. We therefore tested whether we have the power to detect improvements in predictive accuracy between chromosomes, given the observed number of polymorphic sites and expanding k-mer sizes. To do this we created two simulated datasets of substitution counts: one where the true probability of substitution for each k-mer is defined by its 3-mer context, and the other where it is defined by its 9-mer context. We then tested prediction accuracy for the different k-mer models using these simulated datasets to determine whether we would detect an improvement in prediction accuracy.

In the simulated data, the number of substitutions for each 9-mer in each chromosome was determined from a binomial distribution, where the number of trials was equal to the number of each 9-mer, and the probability of success was defined by the observed substitution rates given the 3-mer or 9-mer context. For example, 3-mer and 9-mer probabilities of substitution were calculated from the observed data using all chromosomes, and then these probabilities were used to define the probabilities of success for each trial within chromosomes. Given these simulated datasets, prediction accuracy was tested as mean absolute error calculated from train test resampling between chromosomes as described above.

### Mutational effect and probability of occurrence

We consider the coding consequences for all possible point mutations across all possible codons. In total there are 64 codons and 576 possible point mutations. Of these, 23 cause nonsense mutations, 415 cause missense mutations, and 138 cause synonymous mutations. Each codon point mutation has 256 possible 7-mers and 7-mer substitution rates (supplementary figure 3). We tested the difference in mutation rates between 7-mer substitutions with the potential to cause nonsense, missense, and synonymous mutations with Wilcoxon signed-rank tests. We tested the relationship between the number of potential nonsense, missnese, and synonymous mutations as a function of 7-mer substitution rate using poisson regression. Human 7-mer substitution rates were downloaded from the supplementary information of Aggarwala and Voight (2016). We calculated the Spearman correlation between human and mouse 7-mer substitution rates in three groups: 7-mers substitutions with the potential to cause nonsense mutations, 7-mers substitutions with the potential to cause missense mutations, and 7-mers substitutions with the potential to cause synonymous mutations.

## Author contributions

GP conceived of the idea, conducted the analyses, and wrote the manuscript. HSL, TMMV, AM, CML, and MMS contributed to the manuscript. AM, CML, and MMS helped supervise the project.

