## Supplementary material for "Mechanisms of DNA repair have evolved to minimise the probability of nonsense mutations"

### Supplementary Information

#### Supplementary tables

Supplementary tables 2 to 10 are provided as an excel spreadsheet. Table legends are provided here.

**Supplementary table 1 --** K-mer models under consideration. K-mers are defined by their flanking nucleotides. Note that each k-mer has a complementary k-mer on the reverse strand, effectively halving the total number of k-mers and k-mer changes.

| k-mer model | N possible ancestral k-mers | N possible mutant k-mers | Ancestral k-mer example | | Mutant k-mer example |
|---|---|---|---|---|---|
| 1-mer | 4 | 12 | **C** | > | **A** |
| A-CG | 12 | 18 | **C** G | > | **A** G |
| 3-mer | 64 | 192 | T **C** G | > | T **A** G |
| 5-mer | 1,024 | 3,072 | A T **C** G T | > | A T **A** G T |
| 7-mer | 16,384 | 49,152 | A A T **C** G T G | > | A A T **A** G T G |
| 9-mer | 262,144 | 786,432 | T A A T **C** G T G A | > | T A A T **A** G T G A |

**Supplementary tables 2 to 6 --** 1-mer, 3-mer, 5-mer, 7-mer, and 9-mer counts and probabilities of substitution.

**Supplementary table 7 --** Methylated and unmethylated CG dinucleotide counts and probabilities of substitution.

**Supplementary table 8 --** K-mer counts and interquartile ranges.

**Supplementary table 9 --** 7-mer substitutions with the potential to cause nonsense mutations occur at a significantly lower rate across the neutrally evolving mouse genome than 7-mers substitutions with the potential to cause synonymous mutations. This is consistent for each substitution class.

**Supplementary table 10 --** Rank correlation in human and mouse 7-mer substitution rates. 7-mer substitution rates for substitutions with the potential to cause nonsense mutations are more closely correlated between humans and mice than those with the potential to cause synonymous substitutions.

#### Supplementary Figures

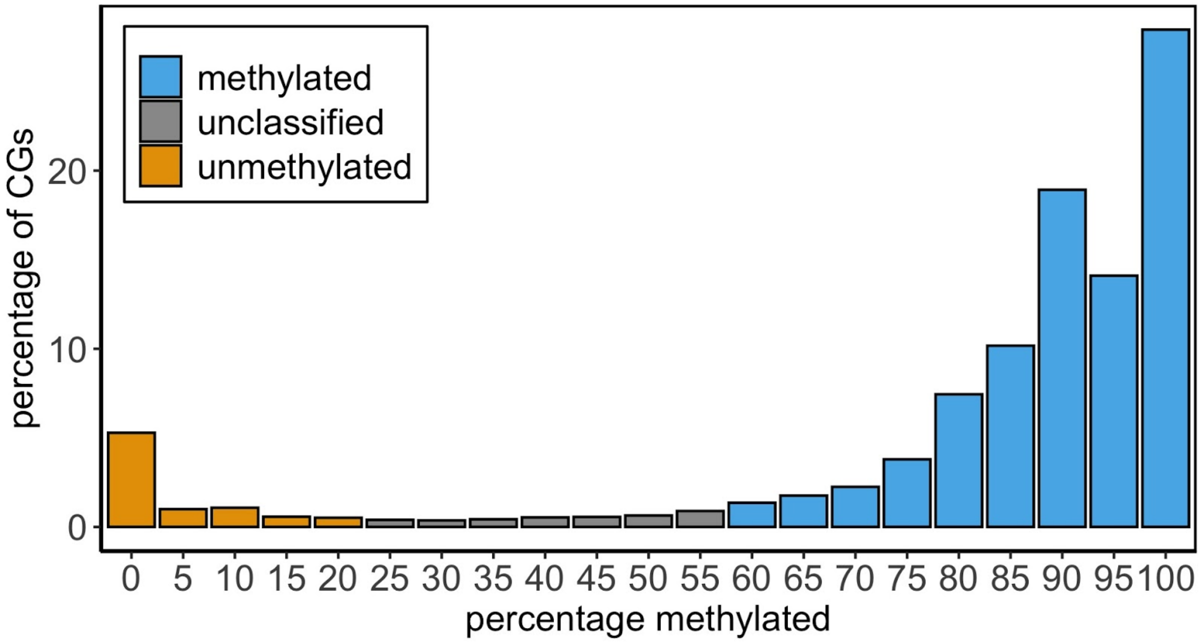


**Supplementary figure 1 --** Distribution of methylation percentage across CG dinucleotides across the mouse genome. We considered autosomal CG dinucleotides with coverage greater than or equal to five (83.4%). Of these, 8.4% have a methylation percentage less than or equal to 20 and are classified as unmethylated, and 87.7% have a methylation percentage more than or equal to 60 and are classified as methylated.

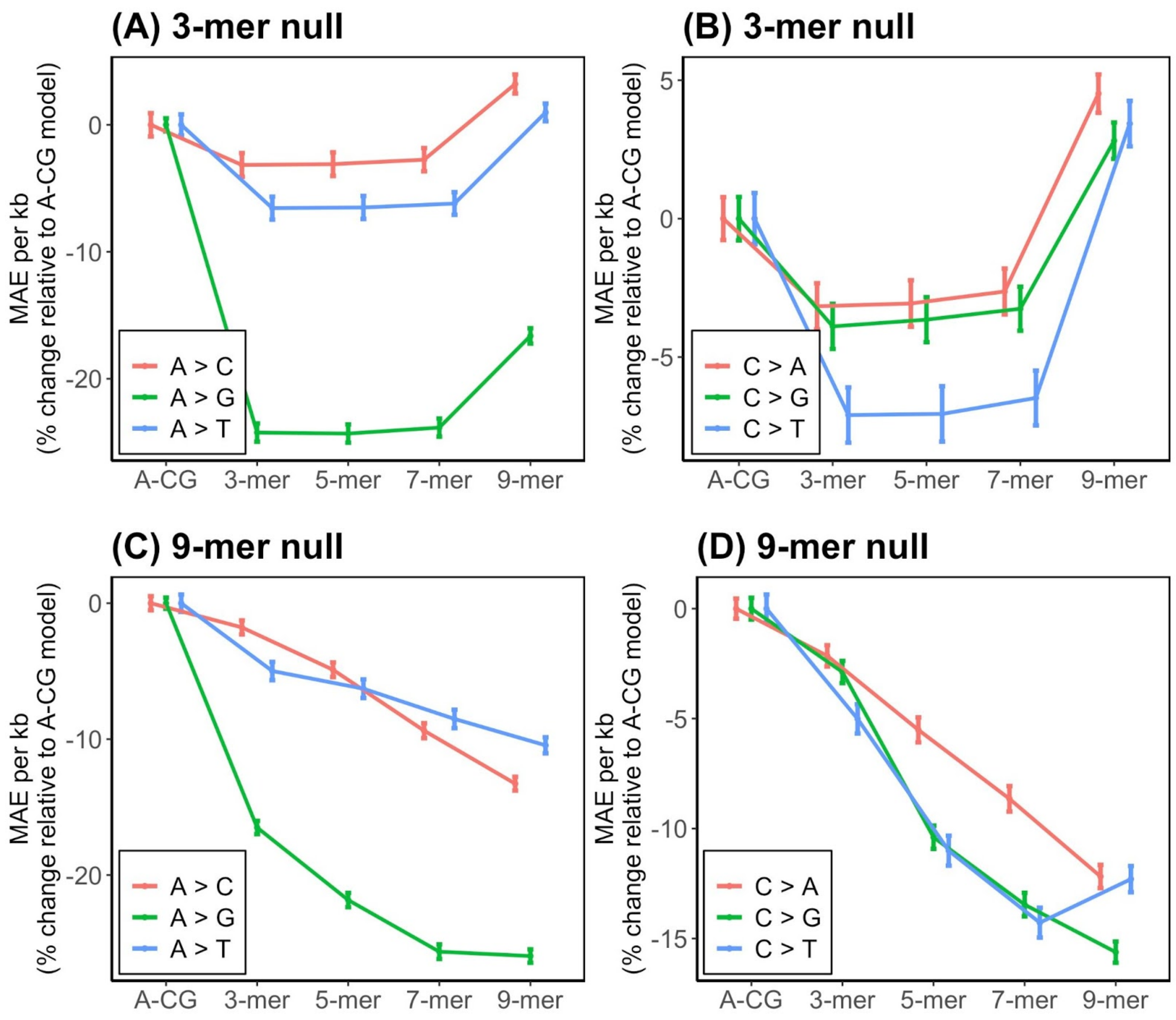


**Supplementary figure 2 --** Mean absolute error (MAE) per kb relative to the A-CG model given simulated probabilities of substitution determined by the 3-mer (A and B) and 9-mer (C and D) context for adenine and cytosine.

Our simulations show that when probabilities of substitution are defined by the 3-mer context, MAE is lower for each substitution class for the 3-mer model compared with the A-CG model. MAE remains consistent for the 5-mer and 7-mer models as the additional bases do not aid prediction. MAE increases, however, for the 9-mer model relative to all other k-mer models. This is likely caused by increased variance in 9-mer probabilities of substitution between chromosomes, resulting from the greater number of parameters and reduced sample size. When probabilities of mutation are defined by their 9-mer context, MAE is lower for each substitution class, except C > T substitutions, for the 9-mer model compared with all other models. This suggests that there is enough power to detect improvements in MAE with the 9-mer model, given the simulated effect size. MAE increases for C > T substitution prediction when considering the 9-mer context compared with the 7-mer.

**ACCAAGA > ACCTAGA (forward strand)**

| | | | | |
|---|---|---|---|---|
| Codon 2 Base 1 | ACC AAG ACC<br>THR LYS THR | ⇨ | ACC TAG ACC<br>THR STOP THR | Nonsense |
| Codon 2 Base 2 | CAC CAA GAC<br>HIS GLN ASP | ⇨ | CAC CTA GAC<br>HIS LEU ASP | Missense |
| Codon 2 Base 3 | CCA CCA AGA<br>PRO PRO ARG | ⇨ | CCA CCT AGA<br>PRO PRO ARG | Synonymous |

**TCTTGGT > TCTAGGT (reverse strand)**

| | | | | |
|---|---|---|---|---|
| Codon 2 Base 3 | GGT CTT GGT<br>GLY LEU GLY | ⇨ | GGT CTA GGT<br>GLY LEU GLY | Synonymous |
| Codon 2 Base 2 | GTC TTG GTG<br>VAL LEU VAL | ⇨ | GTC TAG GTG<br>VAL STOP VAL | Nonsense |
| Codon 2 Base 1 | TCT TGG TGG<br>SER TRP TRP | ⇨ | TCT AGG TGG<br>SER ARG TRP | Missense |

**Supplementary figure 3 --** An example of the coding consequences for A > T substitutions given the ACCAAGA 7-mer context. Each 7-mer substitution can have six different consequences as they have the potential to change each of the three bases in a codon, and this applies to both the forward and reverse strands.

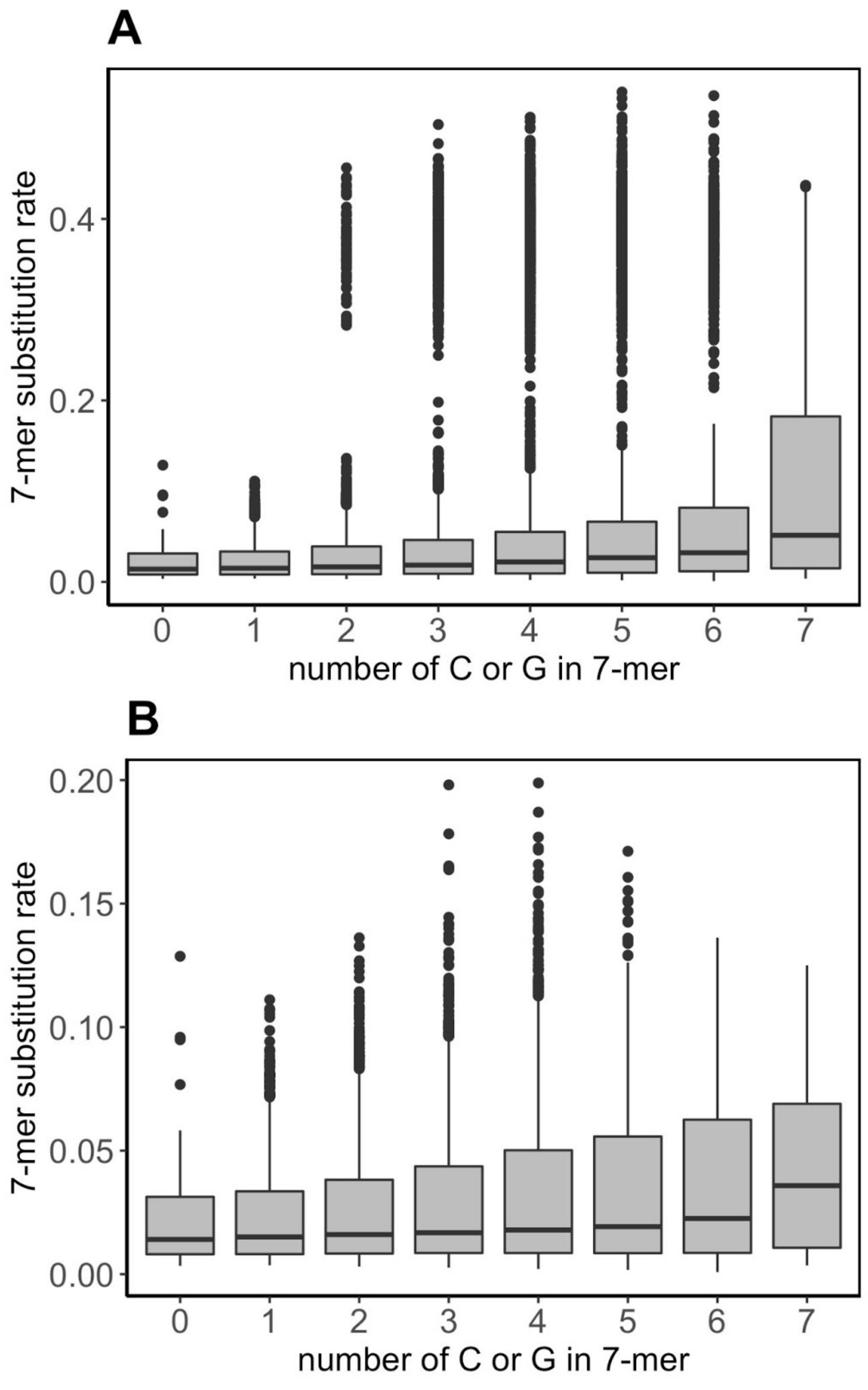


**Supplementary figure 4 --** Substitution rates of 7-mers defined by their composition of cytosine or guanine bases (A). 7-mers with more cytosine or guanine bases have on average higher substitution rates. This relationship persists when CG > TG substitutions are removed from the analysis (B).
